# Intravenous gene transfer throughout the brain of infant Old World primates using AAV

**DOI:** 10.1101/2022.01.08.475342

**Authors:** Miguel R. Chuapoco, Nicholas C. Flytzanis, Nick Goeden, J. Christopher Octeau, Kristina M. Roxas, Ken Y. Chan, Jon Scherrer, Janet Winchester, Roy J. Blackburn, Lillian J. Campos, Cynthia M. Arokiaraj, Timothy F. Miles, Min J. Jang, Julia Vendemiatti, Benjamin E. Deverman, James Pickel, Andrew S. Fox, Viviana Gradinaru

**Author notes:** Authors contributed equally (Miguel R. Chuapoco, Nicholas C. Flytzanis, Nick Goeden).

## Abstract

Adeno-associated viruses (AAVs) can enable robust and safe gene delivery to the mammalian central nervous system (CNS). While the scientific community has developed numerous neurotropic AAV variants for systemic gene-transfer to the rodent brain, there are few AAVs that efficiently access the CNS of higher order primates. We describe here AAV.CAP-Mac, an engineered AAV variant that enables systemic, brain-wide gene delivery in infants of two Old World primate species—the rhesus macaque (*Macaca mulatta*) and the green monkey (*Chlorocebus sabaeus*). We identified CAP-Mac using a multi-species selection strategy, initially screening our library in the adult common marmoset (*Callithrix jacchus*) and narrowing our pool of test-variants for another round of selection in infant macaques. In individual characterization, CAP-Mac robustly transduces human neurons *in vitro* and Old World primate neurons *in vivo*, where it targets all lobes of cortex, the cerebellum, and multiple subcortical regions of disease relevance. We use CAP-Mac for Brainbow-like multicolor labeling of macaque neurons throughout the brain, enabling morphological reconstruction of both medium spiny neurons and cortical pyramidal cells. Because of its broad distribution throughout the brain and high neuronal efficiency in infant Old World primates compared to AAV9, CAP-Mac shows promise for researchers and clinicians alike to unlock novel, noninvasive access to the brain for efficient gene transfer.

## Main Text

Adeno-associated viruses (AAVs) have emerged as dependable and ubiquitous tools for researchers and clinicians alike since their discovery as adenoviral contaminants in the 1960s^1–3^. In the nearly 4 decades following the earliest descriptions of recombinant AAV vectors^4,5^, hundreds of clinical trials serve as a testament that AAVs are nonpathogenic and have the potential to be used safely for long-term expression of genetic payloads^6–9^. There is, however, renewed concern about the safety of high dose systemic AAV following reports of adverse hepatotoxicity^10,11^ and several patient deaths in the clinic^12,13^. The low therapeutic index of systemically administering natural AAV serotypes can require high doses to effectively penetrate tissue, highlighting the need for more efficient, and thus safer, AAVs. In recent years, the field has focused on engineering novel capsids to address this problem, while simultaneously aiming to expand the therapeutic opportunity landscape for gene therapy into disorders not previously approachable with natural AAV serotypes. In parallel, the neuroscience community has also utilized novel AAVs, as several engineered AAV variants can traverse the restrictive blood-brain-barrier (BBB) to systemically deliver genetically-encoded tools to the rodent brain^14–17^, such as GCaMP to detect intracellular calcium gradients^18^. However, the majority of AAV capsid engineering efforts targeting the brain have thus far focused on increasing gene-transfer to the rodent central nervous system (CNS), and direct efforts in non-human primates (NHPs) are sparse. Some recent capsids now enable systemic gene transfer to the brain of the common marmoset^17^ (*Callithrix jacchus*), a New World primate species. But few comparable options exist for Old World primates, which are more evolutionarily related to humans compared to marmosets and are well-established animal models of human cognition, neurodevelopment, neuroanatomy, and physiology^19–21^. To enable research and for greater therapeutic translatability, it is imperative that we advance AAV development for systemic gene transfer to the brains of Old World primates such as the rhesus macaque (*Macaca mulatta*).

Advances in protein engineering, sequencing technologies, and our understanding of AAV structure and function have led to the new neurotropic capsid variants that target the rodent brain. The AAV9 variant AAV-PHP.B was the first variant to unlock efficient widespread gene transfer in adult mammals, traversing the BBB after systemic intravenous (IV) administration in mice^14^. Other variants have since followed with similar or enhanced properties, such as the ability to cross the BBB across different mouse strains, decreased transduction in non-CNS tissue, and biased tropism towards cell-types in the brain^15–17,22,23^. While two groups reported that the BBB-crossing tropism of AAV-PHP.B does not translate to the rhesus macaque^24,25^, studies reporting the systemic translatability of rodent neurotropic capsids in Old World primates are limited. Currently, no AAV enables efficient gene transfer broadly across the Old World primate brain.

In lieu of a capsid variant that can be used for systemic and brain-wide gene transfer in macaques, researchers and clinicians resort to direct injections to circumvent the BBB. However, due to limited spatial distribution, AAVs must typically be administered in multiple locations, invasively penetrating the brain parenchyma each time,^26–33^ with each surgery requiring resource-intensive pre-planning and real-time monitoring of infusions^30–39^. More recently, several groups have utilized intrathecal routes of administration via lumbar puncture (LP)^40^ or intra-cisterna magna (ICM)^41^ injection to overcome the invasiveness and limited spatial distribution of direct injections. However, these intrathecal routes of administration have limited efficacy in the brain^41–45^, and some groups report adverse transduction in non-brain tissue, especially in the dorsal root ganglia^11,45–47^.

Here, we describe AAV.CAP-Mac, an engineered AAV9 variant that efficiently transduces multiple neuronal subtypes throughout cortical and subcortical brain regions of infant Old World primates after IV administration. We initially identified CAP-Mac through 2 rounds of selection in the adult common marmoset (*Callithrix jacchus*). We performed a final round of selection in infant macaques, where CAP-Mac-delivered transgenes were 10- and 6-times more enriched than those delivered by AAV9 in viral DNA and whole RNA brain extracts, respectively. CAP-Mac efficiently transduces the brain in at least two Old World primate species, the rhesus macaque and the green monkey (*Chlorocebus sabaeus*), achieving broader CNS distribution via IV compared to intrathecal administration^45,48^. Furthermore, CAP-Mac targets neuronal cells in the CNS more effectively than its parent AAV9, highlighting the opportunity for broader and more diverse study of the Old World primate brain, as well as the potential for increased therapeutic benefit for disorders affecting neurons. As an example of CAP-Mac’s immediate research utility, we capitalized on its neuronal bias to deliver a cocktail of three fluorescent proteins for Brainbow-like^49,50^, multicolor labeling and morphological tracing in Old World primate brain. By characterizing CAP-Mac in multiple NHP species, we aim to both expand the AAV toolbox available to researchers interested in studying the Old World primate CNS and to highlight the utility of engineering AAVs for increased translatability in higher order mammals.

## Results

### AAV library selection in adult marmosets yields brain-enriched variants

We used a multi-species screening and characterization strategy to select for variants with enhanced BBB-crossing tropism in NHPs (Fig. 1a). To construct the starting capsid library, we inserted 21 degenerate bases ([NNK] x 7) after Q588 in the structural *cap* gene of AAV9, identical to our previously published engineering strategies^14–16^ (Supplementary Fig. 1). We performed 2 rounds of selection in 4 adult male marmosets (2 marmosets per round; 2 × 10^12^ vector genomes [vg] of viral library in each marmoset via IV administration). During round 1 selection, we extracted the brain 4-weeks post-injection and isolated viral DNA from 4 coronal sections per marmoset (8 samples total). Using next-generation sequencing (NGS), we recovered 33,314 unique variants from the 8 samples of brain. We included all recovered variants from round 1 selection plus a codon modified variant of each (66,628 total nucleotide sequences) for round 2 selection.

**Fig. 1:**
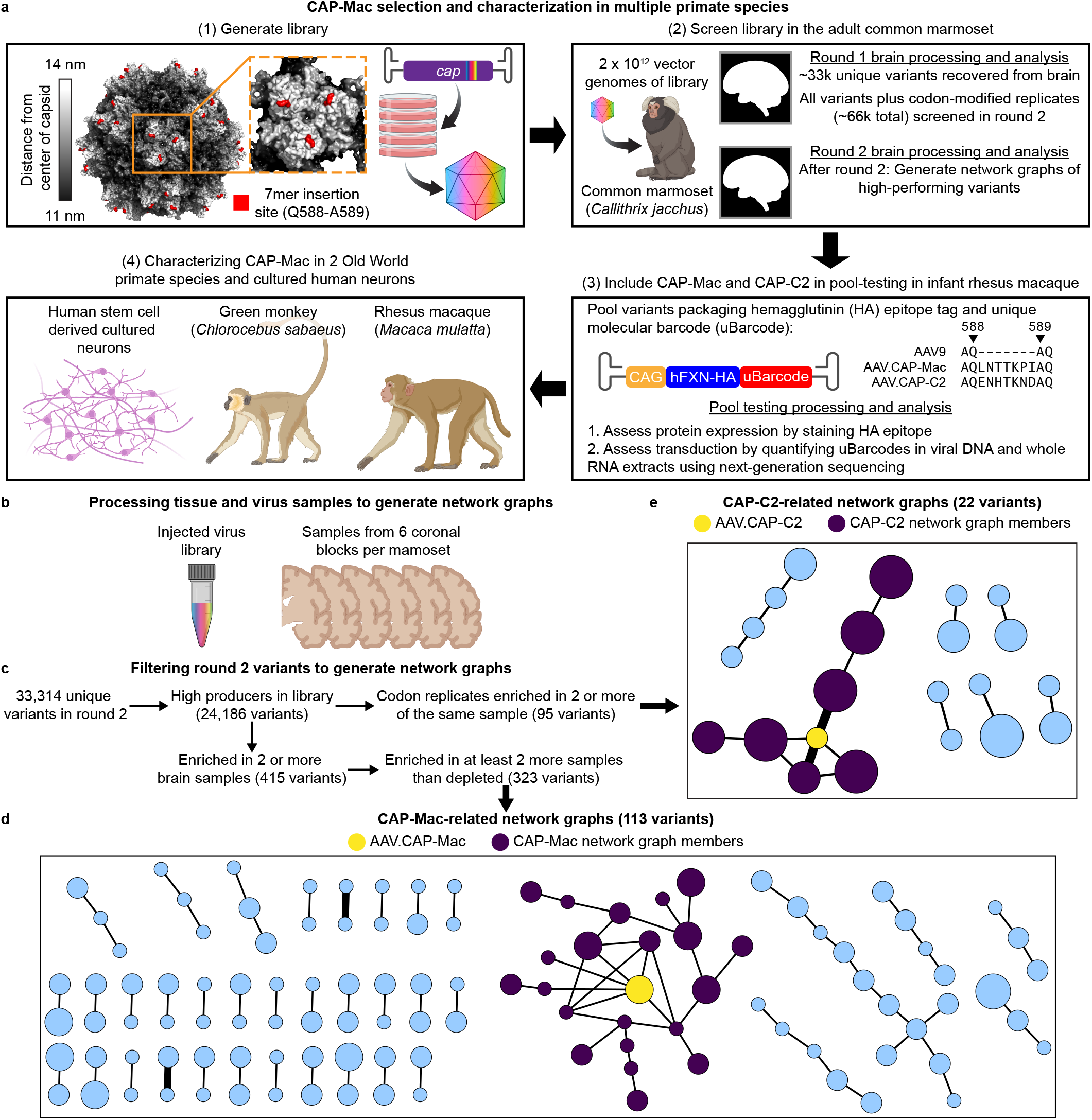
CAP-Mac selection and characterization strategy. **a**, Schematic of the CAP-Mac selection strategy. (1) CAP-Mac is an AAV9 variant that we selected from a library screened in the adult common marmoset. We generated initial diversity by introducing 21 NNK degenerate codons after Q588 in the AAV9 cap gene. (2) We produced and screened the capsid library in 4 adult male marmosets (2 marmosets per round; 2 × 10^12^ vector genomes of library administered intravenously in each marmoset). After the first round of selection, we recovered 33,314 unique amino acid sequences in the brain. For the second round of selection, we generated a synthetic oligo pool containing 66,628 sequences (the ∼33k unique variants plus a codon modified replicate). After the second round of selection, we constructed network graphs of high-performing variants, and selected two capsids—AAV.CAP-Mac and AAV.CAP-C2—to be included in pool selections in infant rhesus macaques. (3) For pool selections, we packaged ssCAG-hFXN-HA with a unique molecular barcode (uBarcode) in the 3’ UTR into 8 different capsids. The construct design enabled us to assess protein expression of the pool by staining for the hemagglutinin (HA) epitope and quantify barcodes in viral DNA and whole RNA extracts. (4) We then individually characterized AAV.CAP-Mac in 2 Old World primate species as well as human cultured neurons. **b**, To generate network graphs, we processed the injected virus library and sampled from each of the 6 brain sections from each animal. **c**, The filtering criteria used to identify high-performing variants. **d, e**, Network graphs for AAV.CAP-Mac (**d**) and AAV.CAP-C2 (**e**). Each node represents a unique variant recoverd from the round 2 selection and each edge represents pairwise reverse Hamming distance ≥ 3.

Notably, in the past we have used Cre-transgenic mouse lines as part of our Cre recombination-based AAV targeted evolution (CREATE) methodology to increase stringency during selections by only recovering variants that undergo cis-Cre-Lox mediated inversion^14,16^. However, since Cre-transgenic marmosets are currently unavailable, we were unable to confer this additional selective pressure during these selections and pursued other strategies to compensate for this loss. We previously demonstrated the utility of clustering capsid variants based on sequence similarity to generate network graphs as an aid in choosing variants for further characterization^16^. Briefly, we filter variants based on user-defined performance criteria and cluster high-performing variants into network graphs, wherein each node is a capsid variant and each edge represents shared sequence identity between related variants (i.e. the pairwise reverse Hamming distance). We reasoned that through this clustering analysis, we could efficiently and productively sample variants from our selections to (1) limit the number of animals used for individual characterization and (2) partially overcome the absence of the selective pressure provided by Cre-transgenic mice in CREATE.

After round 2 selection, we used NGS to quantify the absolute read count of recovered sequences from the injected virus library and from 12 coronal sections (6 sections per marmoset) (Fig. 1b). From the variant read counts, we calculated library enrichment scores, filtered variants, and constructed 2 groups of network graphs (Fig. 1c). One group of network graphs consisted of 113 variants assembled into 37 network graphs, with the largest network containing 22 variants (Fig. 1d). AAV.CAP-Mac (CAP-Mac) is the most interconnected node in this network, sharing an edge with 6 other nodes. A second group of network graphs contained 22 total variants assembled into 7 graphs, with the largest network containing 8 variants (Fig. 1e). AAV.CAP-C2 (CAP-C2), the most interconnected node in this network graph, connects to 4 other nodes. Because of their high connectivity to other variants, we selected CAP-Mac and CAP-C2 for further characterization.

### CAP-Mac is enriched in the infant rhesus macaque CNS compared to other engineered variants

Upon completing library selection in the adult marmoset, we initiated capsid-pool studies in infant rhesus macaques to assess the translatability of several of our engineered AAVs to Old World primates. By pooling several variants into equimolar vector genome doses and administering them to the same rhesus macaque, we could limit the number of animals used for characterization while also assessing variant performance head-to-head, as we have shown that inter-animal variability exceeds intra-animal variability^51^. We pooled a total of 8 capsid variants: CAP-Mac and CAP-C2 as well as the parent capsid, AAV9, plus five other previously engineered AAV controls^15,17^ Each variant packaged a single stranded human frataxin transgene fused to a hemagglutinin (HA) epitope tag under control of the ubiquitous CAG promoter (ssCAG-hFXN-HA) with a unique 12 bp molecular barcode in the 3’ UTR. This construct design allowed us to assess protein expression and localization of the virus pool by staining for the HA tag. We then used NGS to quantify the relative enrichment of each individual barcode in bulk viral DNA and whole RNA extracts from tissue.

We administered 1 × 10^14^ vg/kg of the virus pool (1.25 × 10^13^ vg/kg of each variant) to 2 newborn rhesus macaques via the saphenous vein and extracted brains and livers 4 weeks post-injection. After staining, we observed widespread expression of the HA epitope throughout the brain (Fig. 2a), primarily in cells with neuronal morphology. In cortex and in hippocampus, we observed single cells with clear projections that resemble the apical dendrites of pyramidal cells. Furthermore, we saw increased HA epitope expression in the thalamus and the dorsal striatum (insets).

**Fig. 2:**
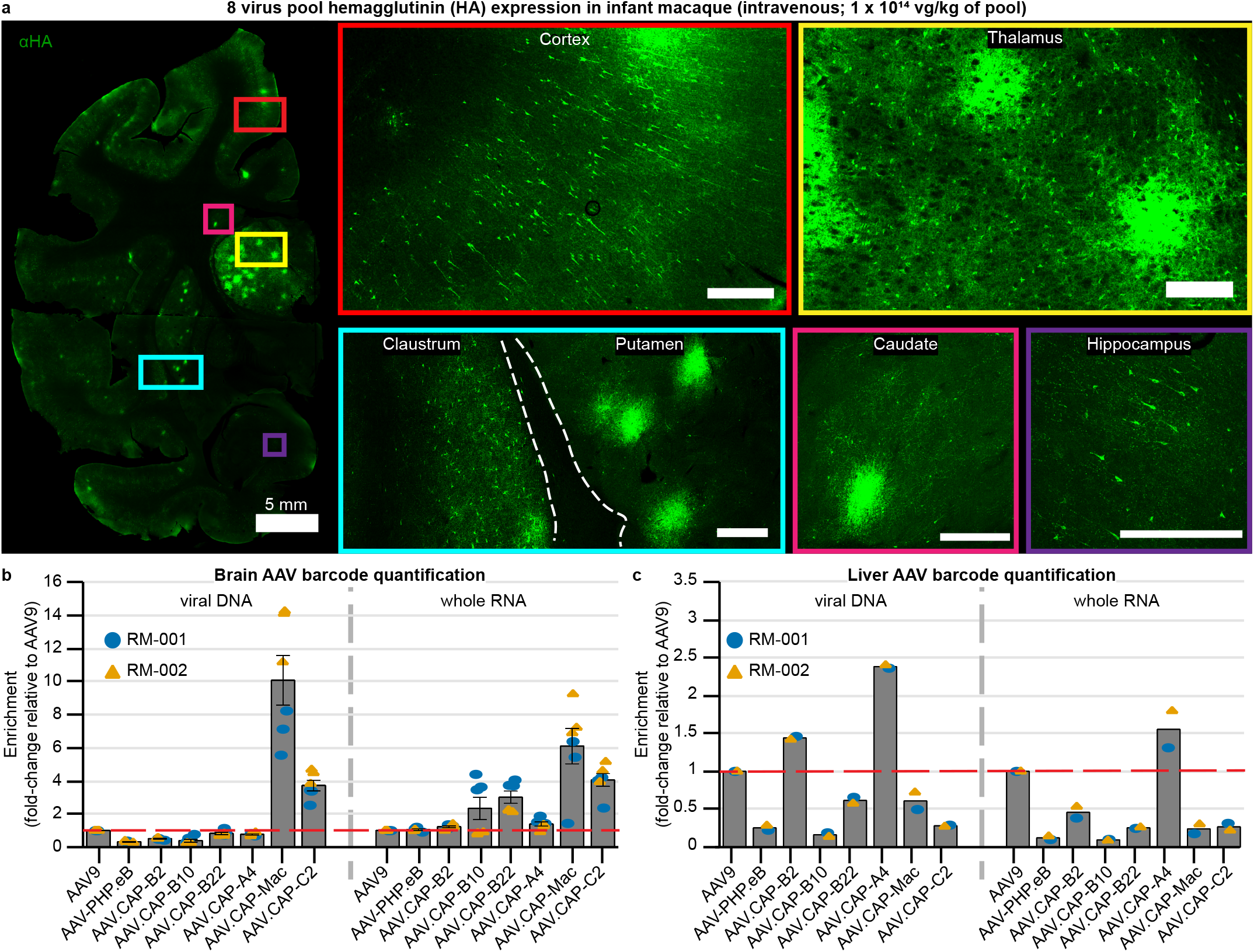
CAP-Mac outperforms other engineered variants in infant rhesus macaque in pool testing. **a**, Representative images of expression in cortex, thalamus, caudate nucleus, putamen, hippocampus and claustrum after intravenous administration of 1 × 10^14^ vg/kg of an 8-capsid pool (1.25 × 10^13^ vg/kg of each variant) packaging hemagglutinin (HA) tagged human frataxin with unique barcode in each capsid. **b, c**, Unique barcode enrichments in viral DNA (left) and whole RNA (right) extracts from the brain (**b**) and the liver (**c**) of two newborn rhesus macaques. Each data point represents the fold-change relative to AAV9 within each sample of tissue. Mean ± s.e.m. shown. The red dotted line denotes AAV9 performance in pool. All scalebars = 500 μm unless otherwise stated.

Given the robust expression of HA epitope, we extracted bulk viral DNA and whole RNA from the brain and liver to quantify the relative enrichment of each barcode. In the brain, AAV.CAP-Mac was the highest performing variant (Fig. 2b). The mean enrichment of the CAP-Mac delivered-barcodes were 10- and 6 -times higher than the AAV9-delivered barcodes in the viral DNA and whole RNA brain extracts, respectively. CAP-C2 barcode mean enrichment was approximately 4-fold higher than AAV9 barcodes in both viral DNA and whole RNA brain extracts. Interestingly in the viral DNA brain extracts, the barcodes of all other variants, which were selected in mice, were on par with AAV9. In the liver, variant barcodes were variably enriched relative to AAV9. Notably, CAP-Mac and CAP-C2 were negatively enriched in the liver, along with some of the previously engineered controls shown to be de-targeted from the liver in rodents^17^ (Fig. 2c). Due to its higher enrichment in the brain, we moved forward with characterizing the single variant CAP-Mac in two species of Old World primates.

### CAP-Mac efficiently transduces the CNS in rhesus macaque and green monkey infants

Due to the apparent neuronal bias of the capsid pool and CAP-Mac’s overabundance in that pool, we explored using CAP-Mac as a research tool to define neuronal morphology with a mixture of fluorescent proteins^15,49,50^. To that end, we intravenously administered a cocktail of 3 CAP-Mac vectors packaging ssCAG-mNeonGreen, ssCAG-mRuby2, and ssCAG-mTurquoise2 in equimolar vector genome ratios into newborn rhesus macaques at a total dose of 5 × 10^13^ vg/kg, via the saphenous vein. We observed widespread expression of all 3 fluorescent proteins in cortex, thalamus (lateral geniculate nucleus), and cerebellum (Fig. 3a-c, respectively). Based on morphology, CAP-Mac primarily transduced neuronal cell types. In the cerebellum and thalamus specifically, we observed a high density of transduced cells, and the highest proportion of co-localization of multiple fluorescent proteins. However, co-localization of 2 or 3 fluorescent proteins appeared to be rare, suggesting that co-infection was uncommon after systemic administration. Fluorescent protein expression appeared to be similar across different coronal slices along the anterior-posterior axis (Fig. 4a). Fluorescent protein expression was robust in all four lobes of cortex and in subcortical areas like the dorsal striatum and hippocampus. We continued to see strong fluorescent protein expression in the thalamus, including in the lateral and medial nuclei, the lateral geniculate nucleus, and the pulvinar. Despite using the ubiquitous CAG promoter, we observed expression almost entirely in neuronal subtypes based on morphology in all regions. Given the broad and robust expression of fluorescent proteins throughout the brain, we were able to assemble morphological reconstructions of both medium spiny neurons (Fig. 4b) and cortical pyramidal cells (Fig. 4c), which are both known to be implicated in human disease^52,53^. In a separate study, we also attempted to administer CAP-Mac via LP administration in infant rhesus macaques, but we report here that efficiency throughout the brain was noticeably decreased compared to the IV administered animals (Supplementary Fig. 2) and especially low in subcortical structures, which has previously been reported^41–45^.

**Fig. 3:**
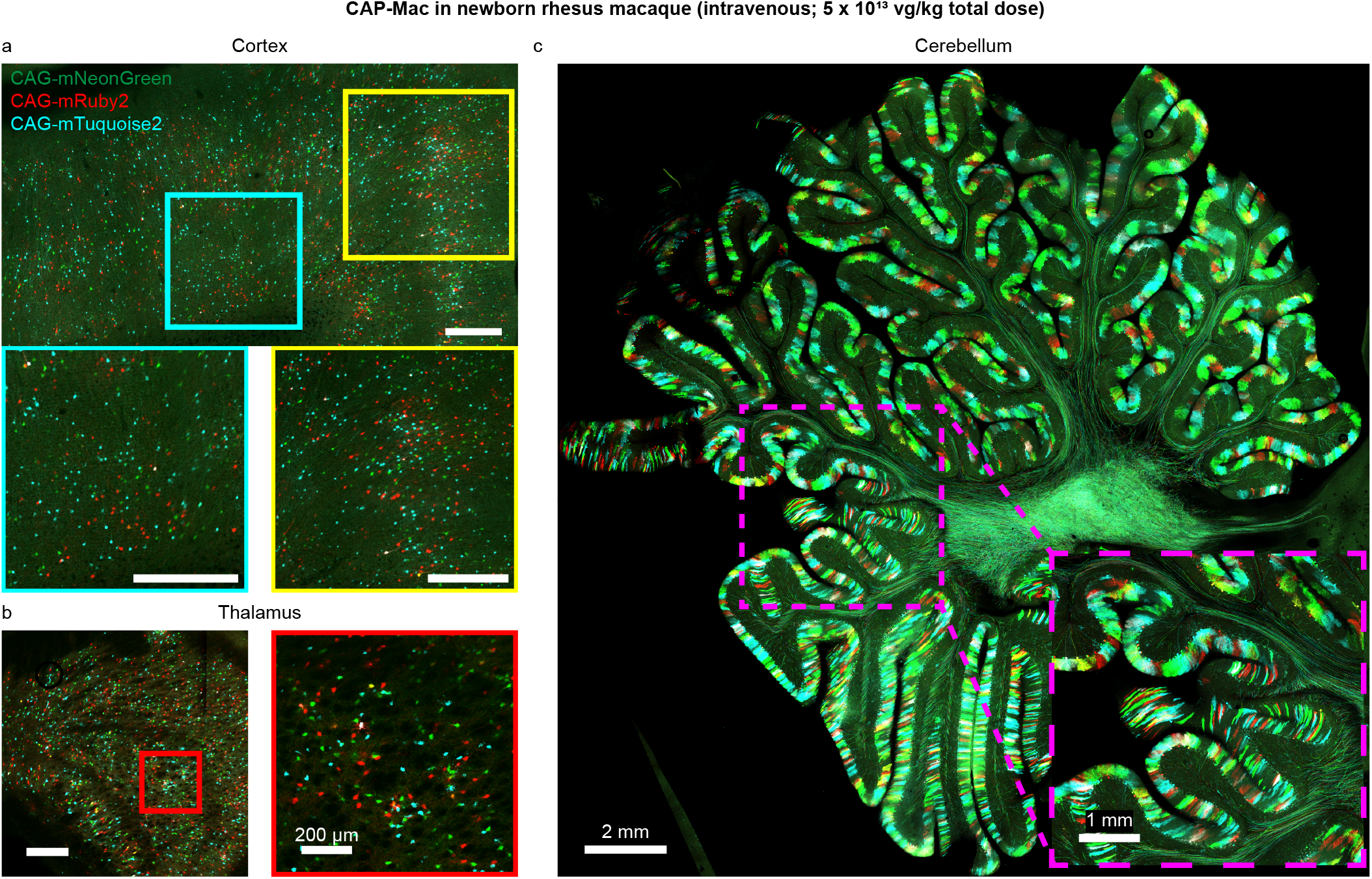
Brain-wide multicolor labeling of neurons in cortex, thalamus, and cerebellum after IV administration of CAP-Mac packaging 3 fluorescent proteins in newborn rhesus macaque. **a, b, c**, Representative images of cortex (**a**), thalamus (lateral geniculate nucleus) (**b**), and cerebellum (**c**) demonstrating widespread expression of a cocktail of 3 fluorescent proteins packaged in CAP-Mac (5 × 10^13^ vg/kg total dose via intravenous administration). All scalebars = 500 μm unless otherwise stated.

**Fig. 4:**
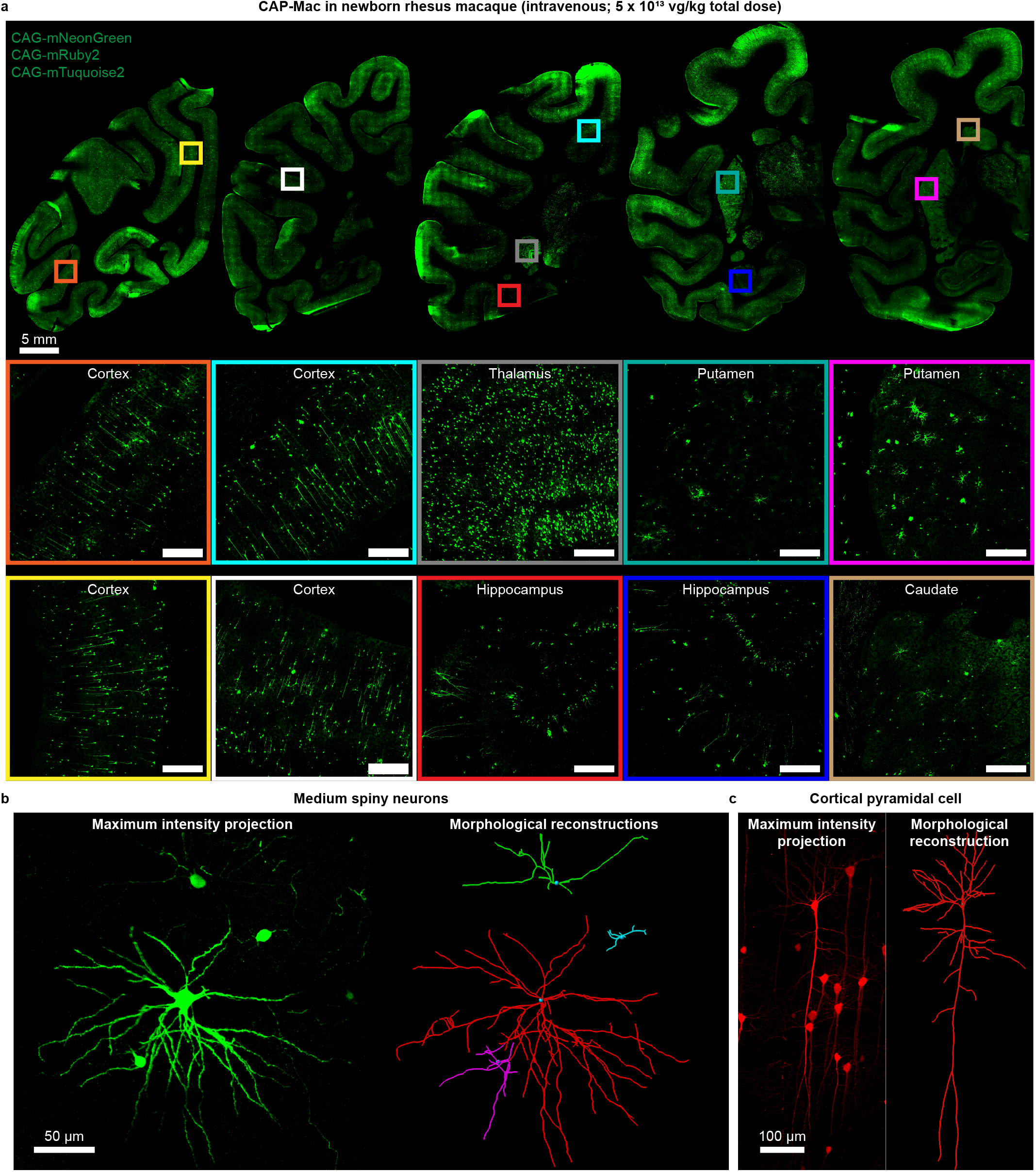
CAP-Mac transduces neurons throughout the newborn rhesus macaque brain, enabling morphological reconstruction of disease-relevant cell types. **a**, Distribution of CAP-Mac expression across coronal slices showing fluorescent protein expression in cortical and subcortical brain regions (insets). Fluorescent proteins are identically pseudocolored. **b, c**, Morphological reconstruction of rhesus macaque medium spiny neurons (**b**) and a cortical pyramidal cell (**c**) in 300 μm sections immersed in RIMS, enabled by intravenous administration of CAP-Mac packaging fluorescent protein. All scalebars = 500 μm unless otherwise stated.

Other engineered AAV variants for BBB-crossing in mice are known to have strain-dependent behavior^16,25,54,55^. Therefore, in parallel to the rhesus macaque experiments, we characterized CAP-Mac in green monkeys, another Old World primate species. We administered AAV9 or CAP-Mac packaging ssCAG-eGFP to individual 8-month-old monkeys at a dose of 7.5 × 10^13^ vg/kg via the saphenous vein. In the CAP-Mac-dosed green monkeys, we saw broad and strong neuronal expression in all four lobes of cortex and in various subcortical regions, including the putamen (Fig. 5a), which is consistent with the expression we observed in the pooled studies (Fig. 2a) and in the rhesus macaque (Fig. 3 and Fig. 4). Notably, we saw particularly strong enhanced green fluorescent protein (eGFP) expression throughout the cerebellum in the CAP-Mac-dosed green monkey. Consistent with other reports, AAV9 tropism appeared to be primarily biased towards cell types with astrocyte-like morphology^41,45,56,57^, and neuronal transduction was low throughout the cortex in AAV9-dosed green monkeys. Conversely, CAP-Mac is biased towards neurons throughout the brain, transducing a higher percentage of neurons than AAV9 in all cortical and subcortical regions that we sampled (Fig. 5b). Viral DNA extracts from various brain regions corroborated this observed increased GFP expression, as recovered eGFP transgene in CAP-Mac green monkeys is consistently higher throughout the brain compared to AAV9-dosed monkeys, suggesting overall higher brain penetrance of CAP-Mac (Fig. 5c and Supplementary Fig. 3a). Interestingly, the cerebellum contained the fewest vector genomes per diploid genome in both CAP-Mac monkeys despite strong eGFP expression, most likely due to the high density of cells and processes within the cerebellum^58,59^. In most non-brain tissue, eGFP biodistribution and expression appeared to be comparable between CAP-Mac and AAV9 treated animals (Supplementary Fig. 3). However, because CAP-Mac transduces a higher percentage of neurons than AAV9, it is worth noting that cell-type tropism differences may continue to persist in non-brain tissue as well, and single-cell studies highlight that even in homogenous cell populations, there is significant viral infection variability^51,60,61^. As such, it is important to consider the role of differential infection dynamics in heterogenous cell types when interpreting viral DNA biodistribution, as measured viral DNA may not correlate to capsid penetrance in tissue in an identical manner across variants and cell types.

**Fig. 5:**
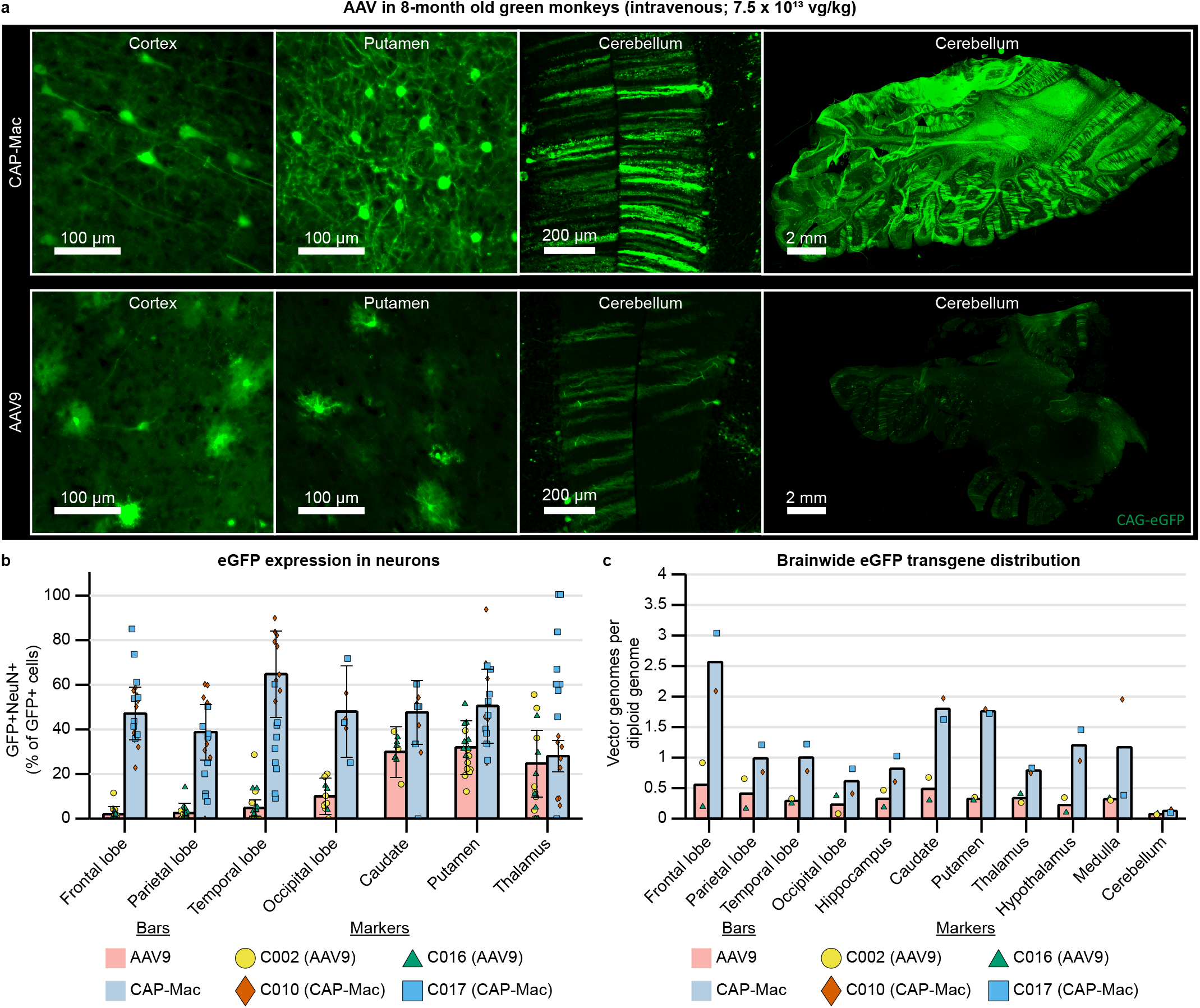
CAP-Mac transduces the green monkey brain more efficiently than AAV9 and is biased towards neurons. **a**, Representative images of various brain regions from green monkeys dosed with CAP-Mac (top) or AAV9 (bottom) packaging ssCAG-eGFP (7.5 × 10^13^ vg/kg via intravenous administration). **b**, NeuN quantification of GFP-transduced cells in images from 7 brain regions of green monkeys. Cells that are positive for both GFP and NeuN are expressed as a percentage of total GFP+ cells in that field of view. Each data point represents quantification of one field of view per condition. **c**, Distribution of CAP-Mac and AAV9-delivered eGFP transgene in 11 brain regions of green monkeys. Each data point represents measured vector genomes per diploid genome in a piece of tissue from each condition. Red bars: mean AAV9 values. Blue bars: mean CAP-Mac values. Yellow circles: measurements from AAV9-treated monkey, C002. Green triangles: measurements from AAV9-treated monkey, C016. Orange diamonds: measurements from CAP-Mac-treated monkey, C010. Blue squares: measurements from CAP-Mac-treated monkey, C017. Mean ± s.e.m. shown (s.e.m. only calculated for samples with n > 2).

Given that we selected CAP-Mac in NHPs and due to its neuronal bias, we wanted to assess the capsid’s utility in rodents. Therefore, we administered CAP-Mac packaging ssCAG-mNeonGreen to C57BL/6J, BALB/cJ, and DBA/2J adult mice via both IV and intracerebroventricular (ICV) administrations. Interestingly, the neuronal bias of CAP-Mac extended to mice when delivered to the adult brain through ICV administration but not IV, where it appeared to primarily transduce cells that make up the vasculature (Supplemental Fig. 4a and b). Furthermore, there is no apparent difference in CAP-Mac tropism across the three mouse strains. We also IV administered CAP-Mac to P0 C57BL/6J mice and observed ssCAG-mNeonGreen expression throughout the brain in various cell types, including neurons, astrocytes, and vasculature (Supplemental Fig. 4c). As rodent experiments are more accessible and have shorter timelines compared to NHPs, the strong neuronal tropism of CAP-Mac in adult mice after ICV administration offers a method to screen genetic cargo in mice prior to applying them in NHPs.

### CAP-Mac strongly transduces human neurons derived from pluripotent stem cells

Given the efficacy of CAP-Mac in penetrating the brain of infant Old World primates and motivated by our observations that CAP-Mac primarily transduces neurons, we wanted to verify whether CAP-Mac offered any improvement over its parent capsid, AAV9, in transducing human neurons. We differentiated cultured induced pluripotent stem cells (iPSCs) derived from humans into mature neurons (Fig. 6a) and incubated cultures with CAP-Mac or AAV9 packaging ssCAG-eGFP across a broad range of doses ranging from 1 vg/cell to 10^6^ vg/cell. We found that GFP expression was noticeably increased in CAP-Mac-administered cultures compared to AAV9-administered cultures (Fig. 6b). Across the aforementioned dose range, we compared the number of transduced cells and eGFP expression per cell between the CAP-Mac and AAV9-treated cultures. AAV9 transduction efficiency achieved an EC_50_=10^4.68^ vg/cell, while CAP-Mac achieved an EC_50_=10^3.03^ vg/cell (Fig. 6c), a 45-fold increase in potency of CAP-Mac in transducing human neurons *in vitro*. Average per cell eGFP expression measured across the population of transduced cells fits a biphasic step function with CAP-Mac reaching the first plateau at a dose roughly two orders of magnitude lower than AAV9 (Fig. 6d). Overall, the increased potency of CAP-Mac in transducing mature human neurons *in vitro* is consistent with the neuronal tropism observed in Old World primates, suggesting a similar mechanism of neuronal transduction across species.

**Fig. 6:**
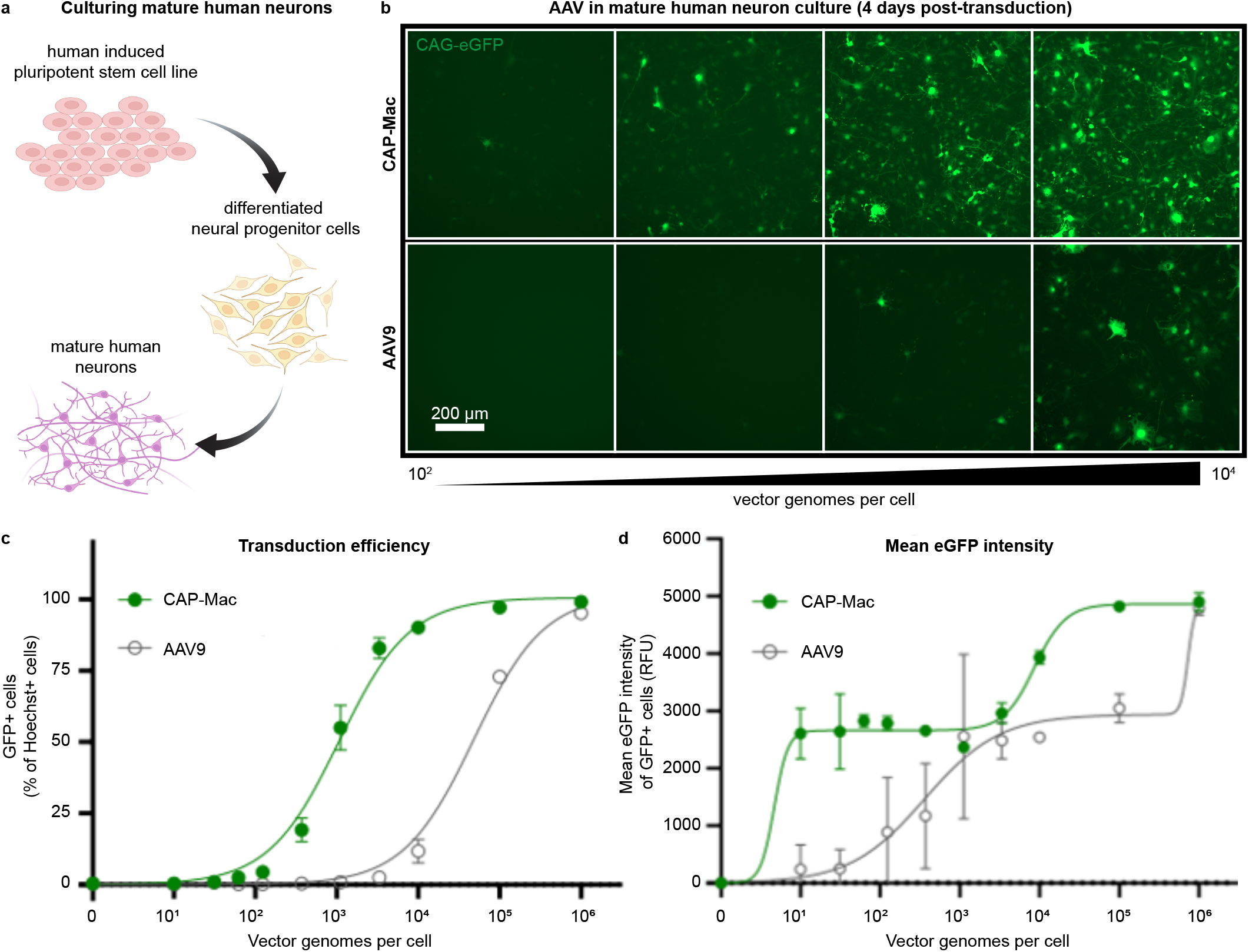
CAP-Mac is more potent at transducing human cultured neurons compared to AAV9. **a**, Differentiation process starting with a human induced pluripotent stem cell line that was differentiated into neural progenitor cells, which were further differentiated into mature neurons. **b**, Representative images of cultured human neurons after 4 days of incubation with either CAP-Mac (top) or AAV9 (bottom) packaging CAG-eGFP across 4 doses of AAV, ranging from 10^2^-10^4^ vector genomes per cell. **c, d**, Dose response curves of AAV9 and CAP-Mac in mature human neuron culture measuring transduction efficiency (**c**) and mean eGFP intensity (**d**).

## Discussion

We describe here the selection and characterization of CAP-Mac, an engineered AAV capsid variant that enables brain-wide transgene expression in Old World primates. CAP-Mac outperforms AAV9 and broadly transduces neurons throughout both cortical and subcortical structures in two Old World primates, the rhesus macaque and the green monkey.

*In vivo* AAV capsid selections have been primarily conducted in mice, partially due to the utility of Cre-transgenic mouse lines to increase selective pressure during selections, as these Cre-based selections can yield neurotropic capsids in as few as two rounds of selection^14,16,17^. However, these engineered variants have thus far failed to translate to NHPs^24,25^. The notable exceptions are AAV.CAP-B10 and AAV.CAP-B22, which were identified using M-CREATE^16^ selections in mice and retain their BBB-crossing and reduced liver tropism in the common marmoset^17^, a New World primate. However, our pool testing experiments here showed that these variants are on par with AAV9 within viral DNA extracts from the brains of infant macaques, an Old World primate. While mice last shared a common ancestor with humans approximately 80-90 million years ago (mya), marmosets and macaques are believed to have shared their last ancestor with humans 35-40 mya and 25-30 mya, respectively^62^. Given this distant shared ancestry, it is therefore not surprising that most variants selected in mice have failed to translate to Old World primates and human patients. Interestingly, we show here in pool studies that variants identified via Cre-independent selections in marmosets and chosen using network graphs (CAP-Mac and CAP-C2), generally outperformed variants identified via Cre-dependent selections in mice (Fig. 2b). This observation suggests that while technological advancements for enhancing selective pressures are important when evolving engineered AAVs *in vivo*, the evolutionary relatedness between the selection-species and target-species is vital to consider if the engineering goal is translatability in higher-order organisms. Notably, several transgenic marmoset lines are currently available^63,64^, and the generation of Cre-transgenic marmosets is currently underway^65^, opening the possibility of performing M-CREATE in NHPs. Interestingly, AAV.CAP-B10 and AAV.CAP-B22 were selected in mice but retain their BBB-crossing tropisms in marmosets. Given that the evolutionary distance between mouse and marmosets (40-55 mya) is slightly larger than the relatedness between marmosets and humans (35-40 mya), this offers hope that NHP selections can identify capsid variants that are efficacious in humans. Overall, CAP-Mac is a testament to the utility of using a selection species that is closely related evolutionarily to the target species.

In the Old World primate studies presented here, we used CAP-Mac exclusively in infants. While this presented several logistical benefits to partially de-risk our characterization efforts (e.g. infants are more likely to be seronegative for neutralizing AAV antibodies and it is more practical to produce sufficient virus to IV dose multiple primates if they weigh less), there is a perception that the developing BBB is “leaky” due to underdeveloped structural integrity, making it more permissive to molecules compared to the adult BBB. However, this interpretation of classical literature has been challenged more recently^66–68^, and contemporary studies report that tight junctions, the main structural unit that enables the BBB to act as a physical barrier between peripheral blood and the extracellular space, are formed embryonically and that the BBB is structurally intact by birth^69–71^. This suggests that CAP-Mac’s brain-penetrating tropism in infant primates and mice is not a passive process, and rather requires active transport across the BBB, most likely via receptor-mediated transcytosis. While the BBB’s structural integrity may be static postnatally, other molecular and cellular developmental processes are dynamic during maturation^69,70,72^, which may dictate tropism differences across developmental states. Such developmental differences in BBB state may be apparent in our studies in mice, where IV administration of CAP-Mac to adult mice results in ssCAG-mNeonGreen expression in vasculature, but the same experiment in P0 mouse pups results in expression in neurons, astrocytes, and vasculature—a stark difference in tropism. The dynamic nature of the molecular and cellular properties of the BBB and the difference in CAP-Mac tropism between adult mice and P0 mouse pups shown here highlight the importance of considering the developmental state when using AAVs both in research and in the clinic.

Given this observation, we are actively working to fully characterize CAP-Mac in infants and adults of various primate species, as a major overarching goal of this study is to define and disseminate a suite of genetic tools to study the NHP brain, especially in Old World primates. This includes characterizing functional cargo that can be paired with CAP-Mac to study the macaque brain. The first such demonstration is included here, where we deliver a cocktail of three fluorescent proteins using CAP-Mac to achieve noninvasive, Brainbow-like^49,50^ labeling in the macaque brain. Because CAP-Mac transduces neurons in cortical and subcortical areas, it provides researchers a more accessible and efficient method to elucidate the morphology of various neuronal cell types, including medium spiny neurons and pyramidal cells. These are both key cell types implicated in neurodegenerative disorders^52,53,73,74^, and the experiments shown here open the door for comparing these cell types in diseased and healthy macaques. Moving forward, the work here will enable major efforts under the NIH BRAIN Initiative^75^, as CAP-Mac-mediated labeling can be readily combined with tissue clearing and imaging techniques to map long-range projections within the Old World primate connectome^76–78^. Such studies present a novel opportunity to study the macaque brain, and more generally, an opportunity to understand the inner workings of the primate CNS.

In addition to CAP-Mac’s utility as a tool to study the primate brain, it may also represent a compelling delivery vehicle for genetic medicine in humans. CAP-Mac, with its efficient neuronal transduction across the macaque brain when delivered IV, provides an unprecedented opportunity to deepen our understanding of the pharmacodynamics of genetic medicines in Old World primate models^32,79,80^. Furthermore, the broad and uniform distribution of CAP-Mac throughout the primate CNS has the potential to provide unprecedented therapeutic access to subcortical and midbrain regions, which has previously proven difficult in NHPs^41–45^. Of further importance for IV delivered gene therapies, the data suggests that CAP-Mac may have reduced tropism towards the liver across primate species. Additionally, CAP-Mac’s enhanced transduction of cultured human neurons supports the potential of CAP-Mac as a clinically relevant gene-delivery vehicle in humans. Overall, the success of the capsid engineering approach we describe here to generate novel variants with BBB-crossing tropism and cell-type bias in Old World primates offers a roadmap for developing the next class of translational gene therapies with improved safety and efficacy profiles.

## Methods

### AAV DNA library generation

We initially generated our diversity at the DNA level, which is used to produce transfection material to produce the AAV capsid library. For the round 1 library, we introduced this genetic diversity using primers containing degenerate nucleotides that were inserted between CAP amino acids 588 and 589^14–16^ (Supplementary Fig. 1a). We used a reverse primer containing 21 degenerate nucleotides ([NNK] x 7) to randomly generate PCR fragments containing unique 7mer sequences inserted into the *cap* genome. For the round 2 DNA library, we used a synthetic oligo pool (Twist Bioscience) as a reverse primer, encoding only variants that we selected for further screening (66,628 DNA oligos total: 33,314 variants recovered after round 1 selections plus a codon-modified replicate of each). All reverse primers contained a 20 bp 5’ overhang complementary with the CAP sequence near the AgeI restriction enzyme sequence and were paired with a forward primer containing a 20 bp 5’ overhang near the XbaI restriction enzyme sequence. We then inserted the PCR fragments containing the diversified region into the rAAV-ΔCAP-in-cis-Lox plasmid via Gibson assembly to generate the resulting AAV DNA library, rAAV-CAP-in-cis-Lox, using NEBuilder HiFi DNA Assembly Master Mix (New England Biolabs, E2621).

### AAV capsid library production

We generated AAV capsid libraries according to previously published protocols^16,81^. Briefly, we transfected HEK293T cells (ATCC, CRL-3216) in 150 mm tissue culture plates using transfection grade, linear polyethylenimine (PEI; Polysciences, Inc). In each plate, we transfected 4 plasmids: (1) the assembled rAAV-Cap-in-cis-Lox AAV DNA library, which is flanked by inverted terminal repeats (ITR) required for AAV encapsidation; (2) AAV2/9 REP-AAP-ΔCAP, which encodes the REP and AAP supplemental proteins required for AAV production with the C-terminus of the CAP gene excised to prevent recombination with the AAV DNA library and subsequent production of replication-competent AAV; (3) pHelper, which encodes the necessary adenoviral proteins required for AAV production; and (4) pUC-18, which contains no mammalian expression vector but is used as filler DNA to achieve the appropriate nitrogen-to-phosphate ratio for optimal PEI transfection. During preparation of the PEI-DNA mixture, we added 10 ng of our AAV DNA library (rAAV-Cap-in-cis-Lox) for every 150 mm dish and combined AAV2/9 REP-AAP-ΔCAP, pUC-18, and pHelper in a 1:1:2 ratio, respectively (40 μg of total DNA per 150 mm dish). At 60 hours post-transfection, we purified AAV capsid library from the both the cell pellet and media using polyethylene glycol precipitation and iodixanol gradient ultracentrifugation. Using quantitative PCR, we then determined the titer of the AAV capsid libraries by amplifying DNaseI resistant viral-genomes relative to a linearized genome standard according to established protocols^81^.

### Marmoset experiments

All marmoset (*Callithrix jacchus*) procedures were performed at the National Institutes of Mental Health (NIMH) and approved by the local Institutional Animal Care and Use Committee (IACUC). Marmosets were born and raised in NIMH colonies and housed in family groups under standard conditions of 27°C and 50% humidity. They were fed ad libitum and received enrichment as part of the primate enrichment program for NHPs at the National Institutes of Health. For all marmosets used in this study, there were no detectible neutralizing antibodies at a 1:5 serum dilution prior to IV infusions (conducted by The Penn Vector Core, University of Pennsylvania). They were then housed individually for several days and acclimated to a new room before injections. Four adult males were used for the library screening, 2 each for first- and second-round libraries. The day before infusion, the animals’ food was removed. Animals were anesthetized with isoflurane in oxygen, the skin over the femoral vein was shaved and sanitized with an isopropanol scrub, and 2 × 10^12^ vg of the AAV capsid library was infused over several minutes. Anesthesia was withdrawn and the animals were monitored until they became active, upon which they were returned to their cages. Activity and behavior were closely monitored over the next 3 days, with daily observations thereafter.

At 4 weeks post-injection, marmosets were euthanized (Euthanasia, VetOne) and perfused with 1X phosphate-buffered saline (PBS). After the round 1 library, the brain was cut into 4 coronal blocks, flash frozen in 2-methylbutane (Sigma Aldrich, M32631), chilled with dry ice, and stored at −80°C for long term storage. After the round 2 library, the brain was cut into 6 coronal blocks, and along with sections of the spinal cord and liver, was flash frozen and stored at −80°C for long term storage.

### Viral library DNA extraction and NGS sample preparation

To extract viral library DNA from marmoset tissue, we previously reported that viral library DNA and endogenous host RNA can be isolated using Trizol by precipitating nucleic acid from the aqueous phase^14,16^. As such, we homogenized 100 mg of spinal cord, liver, and each coronal block of brain in Trizol (Life Technologies, 15596) using a BeadBug (Benchmark Scientific, D1036) and isolated nucleic acids from the aqueous phase according to the manufacturer’s recommended protocol. We treated the reconstituted precipitate with RNase (Invitrogen, AM2288) and digested with SmaI to improve downstream viral DNA recovery via PCR. After digestion, we purified with a Zymo DNA Clean and Concentrator kit (D4033) according to manufacturer’s recommended protocol and stored the purified viral DNA at −20°C.

To append Illumina adapters flanking the diversified region, we first PCR-amplified the region containing our 7mer insertion using 50% of the total extracted viral DNA as a template (25 cycles). After Zymo DNA purification, we diluted samples 1:100 and further amplified around the library variable region with 10 cycles of PCR, appending binding regions for the next PCR reaction. Finally, we appended Illumina flow cell adapters and unique indices using NEBNext Dual Index Primers (New England Biolabs, E7600) via 10 more cycles of PCR. We then gel-purified the final PCR products using a 2% low-melting point agarose gel (ThermoFisher Scientific, 16520050) and recovered the 210 bp band.

For the second-round library only, we also isolated the encapsidated AAV library ssDNA for NGS to calculate library enrichment scores, a quantitative metric that we use to normalize for differences in titer of the various variants in our library (see ref. 11 and the “NGS read alignment and analysis” method section below). To isolate the encapsidated viral genomes, we treated the AAV capsid library with DNaseI and digested capsids using proteinase K. We then purified the ssDNA using phenol: chloroform and amplified viral transgenes by 2 PCR amplification steps to add adapters and indices for Illumina NGS and purified after gel electrophoresis. This viral library DNA, along with the viral DNA extracted from tissue, was sent for deep sequencing using an Illumina HiSeq 2500 system (Millard and Muriel Jacobs Genetics and Genomics Laboratory, Caltech).

### NGS read alignment, analysis, and generating network graphs

Raw fastq files from NGS runs were processed with custom-built scripts (https://github.com/GradinaruLab/protfarm and https://github.com/GradinaruLab/mCREATE)^16^. For the first-round library, we processed the pipeline to process these datasets involved filtering to remove low-quality reads, utilizing a quality score for each sequence, and eliminating bias from PCR-induced mutations or high GC-content. The filtered dataset was then aligned by a perfect string match algorithm and trimmed to improve the alignment quality. We then displayed absolute read counts for each variant during the sequencing run within each tissue, and all 33,314 variants that were found in the brain were chosen for round 2 selections.

After round two selections, we performed the same analysis to display variant absolute read count of the injected virus library and of each variant within each tissue. Additionally, we calculated the library enrichment^16^ for each variant within each tissue:

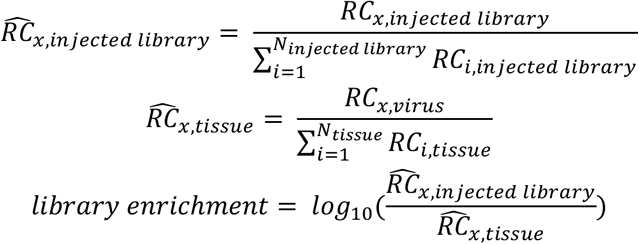

such that for a given sample *y* (e.g. the injected virus library or a tissue sample), *RC*_*x,y*_ is the absolute read count of variant *x, N*_*y*_ is the total number of variants recovered, and 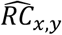 is the normalized read count.

To construct the CAP-Mac sequence clustering graph, we filtered the round 2 NGS data based on the following criteria: (1) ≥ 100 read count in the injected library sample (24,186/33,314 variants), (2) ≥ 0.7 library enrichment score in more than 2 brain samples (415 variants), and (3) at least 2 more brain samples with ≥ 0.7 library enrichment than brain samples with < -0.7 library enrichment (323 variants). To construct the CAP-C2 sequence graph, we filtered the round 2 NGS data based on the following criteria: (1) ≥ 100 read count in the injected library sample and (2) both codon replicates present in at least 2 brain samples with ≥ 0.7 library enrichment (95 variants). These variants were then independently processed to determine pair-wise reverse Hamming distances (https://github.com/GradinaruLab/mCREATE) and clustered using Cytoscape (ver. 3.9.0) as described previously^16^. Networks presented show capsid variants (nodes) connected by edges if the pair-wise reverse Hamming distance is ≥ 3.

### Cloning individual AAV capsid variants

For single variant characterization, we cloned new variant plasmids by digesting a modified version of the pUCmini-iCAP-PHP.eB (Addgene ID: 103005) backbone using MscI and AgeI. We designed a 100 bp primer that contained the desired 21 bp insertion for each capsid variant and the regions complementary to the AAV9 template with ∼20 bp overlapping regions with the digested backbone. We then assembled the variant plasmid using NEBuilder HiFi DNA Assembly Master Mix, combining 5 μL of 200 nM primer with 30 ng of digested backbone in the reaction mixture.

### Individual AAV production and purification

To produce variants for pool testing, we followed our previously published protocol^81^ using 150 mm tissue culture dishes. For individual AAV.CAP-Mac and AAV9 characterization *in vivo* and *in vitro*, we adopted our published protocol to utilize ten-layer CellSTACKs (Corning, 3320) to efficiently produce viruses at high titer to dose rhesus macaques and green monkeys. Specifically, we passaged 20 150-mm dishes at approximately 70% confluency into a 10-layer CellSTACK 24 h before transfection. On the day of transfection, we prepared the DNA-PEI transfection mixture for 40 150-mm dishes and combined the transfection mixture with media and performed a complete media change for the CellSTACK. We collected and changed media at 72 h post-transfection similarly to production in 150 mm dishes. At 120 h post-transfection, we added ethylenediaminetetraacetic acid (EDTA, Invitrogen, 15575020) to a final concentration of 10 mM and incubated at 37°C for 20 min, occasionally swirling and tapping the sides of the CellSTACK to detach the cells. We then removed the media and cell mixture and proceeded with the AAV purification protocol^81^. Of note, during the buffer exchange step after ultracentifugation, we used centrifugal protein concentrators with polyethersulfone membranes (Thermo Scientific, 88533) instead of Amicon filtration devices and used Dulbecco’s PBS supplemented with 0.001% Pluronic^®^ F-68 (Gibco, 24040032).

### Rodent experiments

All rodent procedures were performed at California Institute of Technology (Caltech) and were approved by the local IACUC. We purchased C57BL/6J (000664), BALB/cJ (000651), and DBA/2J (000671) mice (all males, 6–8 weeks old) from The Jackson Laboratory. For IV administration in mice, we delivered virus through the retro-orbital sinus^81,82^ using a 31 G insulin syringe (BD, 328438). For intracerebroventricular administration in mice, we injected into the lateral ventricle. Briefly, we anesthetized mice using isoflurane (5% for induction, 1-3% for maintenance) with 95% O_2_/5% CO_2_ (1 L/min) and mice were head-fixed in a stereotaxic frame. After shaving the head and sterilizing the area with chlorohexidine, we administered 0.05 mL of 2.5 mg/mL bupivacaine subcutaneously, and a midline incision was made and the skull was cleaned of blood and connective tissue. After leveling the head, burr holes were drilled above the lateral ventricles bilaterally (0.6 mm posterior to bregma, 1.15 mm from the midline). Viral vectors were aspirated into 10 μL NanoFil syringes (World Precision Instruments) using a 33-guage microinjection needle, and the needle was slowly lowered into the lateral ventricle (1.6 mm from the pial surface). The needle was allowed to sit in place for approximately 5 min and 3-5 μL of viral vector was injected using a microsyringe pump (World Precision Instruments, UMP3) and pump controller (World Precision Instruments, Mircro3) at a rate of 300 nL/min. All mice received 1 mg/kg of buprenorphine SR and 5 mg/kg of ketoprofen subcutaneously intraoperatively and 30 mg/kg of ibuprofen and 60 mg/kg of Trimethoprim/Sulfamethoxazole (TMPS) for 5 days post-surgery. After 3 weeks of expression, all mice were perfused with PBS and fixed in 4% paraformaldehyde (PFA). All organs were extracted, incubated in 4% PFA overnight, transferred into PBS supplemented with 0.01% sodium azide, and stored at 4°C for long-term storage. We sliced the brain into 100 μm sections by vibratome (Leica Biosystems, VT1200S), mounted in Prolong Diamond Antifade (Invitrogen, P36970), and imaged using a confocal microscope (Zeiss, LSM 880).

### Rhesus macaque experiments

All rhesus macaque (*Macaca mulatta)* procedures were performed at the California National Primate Research Center (CNPRC) at UC Davis and were approved by the local IACUC. Infant macaques were weaned at birth. Within the first month, macaques were infused with AAV vectors either intravenously (IV) or intrathecally. For IV injections, animals were anesthetized with ketamine (0.1 mL) and the skin over the saphenous vein was shaved and sanitized. AAV (see Supplementary tables 1 and 2) was slowly infused into the saphenous vein over ∼1 min in < 0.75 mL of phosphate buffered saline. For IT injections, animals were administered a sedative intramuscularly and the area of skin at the neck was shaved and aseptically prepared. A needle was advanced into the cisterna magna to remove a small amount of CSF proportional to the amount of fluid injected. Then, a sterile syringe containing the sterile preparation of the AAV proportional to the amount of fluid collected was aseptically attached and slowly injected. All animals were monitored during recovery from sedation, throughout the day, and then daily for any adverse findings. All monkeys were individually housed within sight and sound of conspecifics. Tissue was collected 4-11 weeks after injection. Animals were deeply anesthetized and received sodium pentobarbital in accordance with guidelines for humane euthanasia of animals at the CNPRC. All material injected into rhesus macaques were free of endotoxins (<0.1 EU/mL), and protein purity was confirmed by sodium dodecyl sulphate–polyacrylamide gel electrophoresis (SDS-PAGE). See Supplementary tables 1 and 2 for route of administration, AAV variants, viral dose, genetic cargo, and duration of expression for each experiment.

#### Pool testing in infants

Macaques were perfused with ice cold RNase-free PBS. At the time of perfusion, one hemisphere of brain was flash-frozen and the other hemisphere was sectioned into 4 mm coronal blocks and post fixed in 4% PFA for 48 hours and transferred to Caltech for further processing. For HA staining, we incubated slices with rabbit anti-HA (1:200; Cell Signaling Technology, 3724), performed 3-5 washes with PBS, incubated with donkey anti-rabbit IgG (1:200; Jackson ImmunoResearch, 711-605-152), and washed 3-5 times before mounting. We diluted all antibodies and performed all incubations using PBS supplemented with 0.1% Triton X-100 (Sigma-Aldrich, T8787) and 10% normal donkey serum (Jackson ImmunoResearch, 017-000-121) overnight at room temperature with shaking.

To isolate viral DNA and whole RNA, 100mg slices from brain and liver were homogenized in Trizol (Life Technologies, 15596) using a BeadBug (Benchmark Scientific, D1036) and total DNA and RNA were recovered according to the manufacturer’s recommended protocol. Recovered DNA was treated with RNase, underwent restriction digestion with SmaI, and purified with a Zymo DNA Clean and Concentrator Kit (D4033). Recovered RNA was treated with DNase, and cDNA was generated from the mRNA using Superscript III (Thermo Fisher Scientific, 18080093) and oligo(dT) primers according to the manufacturer’s recommended protocol. We used PCR to amplify the barcoded using 50ng of viral DNA or cDNA as template. After Zymo DNA purification, we diluted samples 1:100 and further amplified around the barcode region using primers to append adapters for Illumina next-generation sequencing. After cleanup, these products were further amplified using NEBNext Dual Index Primers for Illumina sequencing (New England Biolabs, E7600) for ten cycles. We then gel-purified the final PCR products using a 2% low-melting point agarose gel (ThermoFisher Scientific, 16520050). Pool testing enrichment was identically to library enrichment, but is represented in Fig 2b and c on a linear scale.

#### Individual characterization of CAP-Mac in infants

Macaques were perfused with PBS and 4% PFA. The brain was sectioned into 4 mm coronal blocks and all tissue was post-fixed in 4% PFA for 3 days before storage in PBS. All tissue was transferred to Caltech for further processing. Brains and liver were sectioned into 100 μm slices using vibratome. Sections of spinal cord were incubated in 30% sucrose overnight and embedded in Optimal Cutting Temperature Compound (Scigen, 4586) and sectioned into 50 μm slices using cryostat (Leica Biosystems, CM1950). All slices were mounted using Prolong Diamond Antifade and imaged using a confocal microscope. For GFP staining of brain slices from the LP administered macaque, we incubated slices with chicken anti-GFP (1:500; Aves Bio, GFP-1020), performed 3-5 washes with PBS, incubated with donkey anti-chicken IgY (1:200; Jackson ImmunoResearch, 703-605-155), and washed 3-5 times before mounting. We diluted all antibodies and performed all incubations using PBS supplemented with 0.1% Triton X-100 (Sigma-Aldrich, T8787) and 10% normal donkey serum (Jackson ImmunoResearch, 017-000-121) overnight at room temperature with shaking.

For morphological reconstruction, we sectioned brain into 300 μm sections and incubated them in refractive index matching solution (RIMS)^83^ for 72 hours before mounting on a slide immersed in RIMS. We imaged using a confocal microscope and 25x objective (LD LCI Plan-Apochromat 25x/0.8 Imm Corr DIC) using 100% glycerol as the immersion fluid. We captured tiled, Z-stacks (1024×1024 each frame using suggested capture settings) around cells of interest, and cropped appropriate fields of view for tracing. Tracing was done Imaris using the semi-automated and automated methods.

### Green monkey experiments

All green monkey (*Chlorocebus sabaeus*) procedures were performed at Virscio, Inc. and approved by their IACUC. All monkeys were screened for neutralizing antibodies and confirmed to have < 1:5 titer. At approximately 7-8 months of age, monkeys were dosed intravenously. Dose formulations were allowed to equilibrate to approximately room temperature for at least 10 minutes, but no more than 60 minutes prior to dosing. IV dose volumes were based on Day 0 body weights. Animals were sedated with ketamine (8 mg/kg) and xylazine (1.6 mg/kg). The injection area was shaved and prepped with chlorohexdrine and 70% isopropyl alcohol, surgically scrubbed prior to insertion of the intravenous catheter. Dosing occurred with a single intravenous infusion on Day 0 via a saphenous vein administered using a hand-held infusion device at a target rate of 1 mL/minute. General wellbeing was confirmed twice daily by cage side observation beginning one week prior to dosing. At the scheduled sacrifice time, monkeys were sedated with ketamine (8-10 mg/kg IM) and euthanized with sodium pentobarbital (100 mg/kg IV to effect). Upon loss of corneal reflex, a transcardial perfusion (left ventricle) was performed with chilled phosphate buffered saline (PBS) using a peristaltic pump set at a rate of approximately 100 mL/min until the escaping fluid ran clear prior to tissue collection. Cubes of tissue were collected from the left brain hemisphere and various other organs and frozen in the vapor phase of liquid nitrogen for further processing for biodistribution. The right brain hemisphere was removed and cut into ∼4 mm coronal slices and post-fixed intact with approximately 20 volumes of 10% neutral-buffered formalin (NBF) for approximately 24 hours at room temperature.

Genomic DNA was extracted from CNS and peripheral tissues using the ThermoFisher MagMax DNA Ultra 2.0 extraction kit (Catalog number: A36570). DNA was assessed for yield by fluorometric quantification with the Qubit dsDNA assay. Approximately 20 ng of DNA was loaded into each 20 μL reaction and plates were run on the BioRad CFX Connect Real-Time PCR Detection System (Catalog number: 1855201). The viral copy number assay was validated for specificity by detection of a single amplified product, sensitivity by assessing the lower limit of detection to be greater than 10 copies per reaction and linearity by ensuring the standard curve r^2^ was > 0.95. Reactions were assembled in FastStart Universal SYBR Green Master (Rox) (catalogue number: 4913850001). The following were the sequences of the primers: forward ACGACTTCTTCAAGTCCGCC, reverse TCTTGTAGTTGCCGTCGTCC.The following PCR protocol was used an initial denaturation step of 95 °C for 180 seconds, followed by 40 cycles of 95 °C for 15 seconds, 60 °C for 60 seconds; with an imaging step following each 60 °C cycle. Standard curve generated with linearized plasmid containing the GFP template sequence present in the virus from 1e8-1e0 copies, diluted in naïve untreated macaque DNA samples prepared using an identical kit to the samples in this study to control for matrix effects. Copies of viral DNA were calculated from the standard curve using the equation for the line of the best fit. MOI values were calculated based on the measured total genomic weight of the host cell DNA per reaction.

Post fixation, tissues were placed into 10% > 20% > 30% sucrose for 24 hours each at 4 °C then embedded in Optimal Cutting Temperature Compound and stored at -80 °C until cryosectioning. Tissue blocks were brought up to -20 °C in cryostat before sectioning into 30 μm slices and dry-mounted onto slides after cryosectioning. After sectioning, the slides were left at room temperature overnight to dry. To assist in neuron quantification, we stained sections with the following antibodies and concentrations: rabbit anti-GFP (1:100; Millipore-Sigma, AB3080) and mouse anti-NeuN (1:500; Millipore-Sigma, MAB377). For secondary antibody staining, the following secondary antibodies and concentrations were used: donkey anti-rabbit Alexa Fluor 488 (1:500; Invitrogen, A21206) and donkey anti-mouse Alexa Fluor 647 (1:500; Invitrogen, A31571). All antibodies were diluted with 1X PBS supplemented with 0.25% Triton X-100 (PBST) and 5% normal donkey serum. Primary antibody incubations were left overnight at room temperature. Sections were then washed with PBST. Secondary antibody incubations were 2 hours at room temperature. The sections were washed 3x in PBST. Sections were incubated in DAPI solution (1:10,000; Invitrogen, D1306) at room temperature for 5 minutes, then washed. Sections were coverslipped using Prolong Diamond Antifade.

3 sections per animal were stained and imaged. Each section was imaged in triplicate with each ROI having a total of 9 images. Tissue ROIs were imaged with a Keyence BZ-X800 with the following acquisition parameters: GFP (1/500 s), Cy5 (1 s), DAPI (1/12 s), High Resolution, Z-stack @ 1.2 um pitch. The following brain subregions were imaged frontal, parietal, temporal, occipital cortices, cerebellum, caudate, putamen, and thalamus (medial, ventral lateral, and ventral posterior nuclei). A semi-automated cell counting method was performed via ImageJ for quantification. Using thresholds and particle analysis, we were able to quantify NeuN positive and DAPI positive cells. Using ImageJ’s cell counter, we manually counted GFP positive and GFP & NeuN double-positive cells.

### Induced pluripotent stem cell (iPSC) experiments

Neuronal cultures were produced by differentiating and maturing iPSC-derived neural progenitor cells with Stemdiff™ Forebrain Differentiation and Maturation kits (StemCell #08600, #08605 respectively), according to their vendor protocols. Neural progenitor cells were produced by differentiation of the foreskin fibroblast-derived iPSC line: ACS™-1019 (ATCC# DYS-0100), with Stemdiff™ SMADi Neural Induction kits (StemCell l#08581), selection with Stemdiff™ Neural Rosette Selection Reagent (StemCell l#05832), and expansion in Stemdiff™ Neural Progenitor Media (StemCell l#05833), according to their vendor protocols. Neurons were matured a minimum of 8 days prior to replating for transduction.

Mature neuronal cultures, seeded 15,000 cells/well in polyornithine and laminin coated, black-walled 96 well optical plates, were cultured an additional 4 days prior to transduction. Replicate wells were transduced with virus serially diluted across six orders of magnitude in 90% maturation media and 10% OptiproSFM. 4 days post-transduction, cultures were fixed with 4% paraformaldehyde and counterstained with 1 ug/ml Hoechst 33322. Identification of transduced cells was determined by imaging 60 fields/well, using two channel fluorescence detection (Hoechst at ex386/em440, eGFP ex485/em521) on a CellInsight CX5 HCS Platform. Individual cells were identified by Hoechst detection of their nuclei and applying size and contact constrained ring masks to each cell. Cell transduction was determined by measuring an eGFP fluorescence above a threshold level within an individual ring mask. For each population, the percentage of transduced cells was plotted vs the applied dose. Curve-fits and EC_50_ values were determined with a Prism GraphPad [agonist] vs response (three parameter) regression method. To report per cell eGFP expression efficiencies, the eGFP spot fluorescence intensities were averaged from each ring mask across a minimum of 5000 cells/well. Curve fits were obtained using Prism GraphPad Biphasic, X as concentration regression method.

## Acknowledgements

We wish to thank the entire Gradinaru laboratory and members of Capsida Biotherapeutics for helpful discussions. We thank Deborah Lidgate for her helpful input in writing and planning the paper. We thank Catherine Oikonomou for help with manuscript editing. We thank Máté Borsos for his assistance in mouse pup injections. We thank Xinhong Chen, Anat Kahan, and Gerard M. Coughlin for their helpful discussion and input in planning rhesus macaque experiments and experimental cargo. Capsida would like to thank Michael Weed and the whole team at Virscio, Inc. for their help with the design and execution of green monkey experiments. We are grateful to Igor Antoshechkin and the Millar and Muriel Jacobs Genetics and Genomics Core at the California Institute of Technology for assistance with next-generation sequencing. We are grateful to the research and veterinarian staff at the California National Primate Research Center (CNPRC) for their aid with studies in rhesus macaques. This work was funded by grants from the National Institutes of Health (NIH) to V.G. (NIH Pioneer DP1OD025535), to the California National Primate Research Center (NIH: P51OD011107), and BRAIN Armamentarium U01 UMH128336A (to V.G., T.F.M., and A.S.F.). Additional funding includes: (to V.G. and A.S.F.) The Michael J. Fox Foundation for Parkinson’s Research as part of the Aligning Science Across Parkinson’s Initiative (ASAP-020495). Figures were created using images from BioRender.com.

## Author contributions

M.R.C. and N.C.F. wrote the manuscript, with input from all authors. M.R.C. designed, performed, and analyzed the data for the rhesus macaque and rodent experiments and prepared all figures. N.C.F., K.C. and B.E.D. designed, N.C.F. and K.C. performed, and N.C.F. analyzed the associated data for the viral library screening experiments in common marmosets. N.C.F. and N.G. designed, performed and analyzed the data of the pooled testing experiments in rhesus macaques. N.C.F. and N.G. designed, N.C.F. N.G., J.C.O. and K.M.R. performed the green monkey experiments, and J.C.O. and K.M.R. analyzed the associated data and helped prepare the associated figures. J.S., J.W. and R.J.B. designed and performed the human neuron experiments, analyzed the associated data and prepared the associated figures. L.J.C. designed and performed the rhesus macaque experiments. C.M.A. performed the rhesus macaque spinal cord and dorsal root ganglia analysis and imaging. T.F.M. analyzed the round 2 viral library screening experiment in the common marmoset and generated the sequence clustering graphs. M.J.J. helped with the imaging analysis. J.V. helped perform the rhesus macaque neuron tracing. J.P. supervised aspects of the common marmoset experiments. A.S.F. designed, performed, and supervised aspects of the rhesus macaque experiments. N.C.F. supervised all aspects of the green monkey and iPSC work. V.G. supervised all aspects of the library screening, pooled testing and rhesus macaque work and contributed to associated experimental design, data analysis, and manuscript writing.

