## Supplemental tables and figures for "Intravenous gene transfer throughout the brain of infant Old World primates using AAV"

**Supplementary table 1: Rhesus macaque information for variant pool testing.**

| Macaque ID | Age (days) | Sex | Route of administration | Weight at injection (kg) | Total dose (vg/kg) | Expression length (days) |
|---|---|---|---|---|---|---|
| RM-001 | 1 | Male | Intravenous | 0.602 | $1 \times 10^{14}$ | 31 |
| RM-002 | 1 | Male | Intravenous | 0.454 | $1 \times 10^{14}$ | 31 |

**All macaques in Supplementary table 1 were injected with pool of 8 variants (AAV9, PHP.eB, CAP-A4, CAP-B2, CAP-B10, CAP-B22, CAP-C1, CAP-C2) packaging ssCAG-hFXN-HA-uBarcode.*

**Supplementary table 2: Rhesus macaque information for individual characterization of CAP-Mac.**

| Macaque ID | Age (days) | Sex | Route of administration | Cargo | Weight at injection (kg) | Total dose (vg/kg) | Expression length (days) |
|---|---|---|---|---|---|---|---|
| RM-008 | 29 | Male | Intrathecal-lumbar puncture | CAG-GCaMP7s | 0.74 | $2.5 \times 10^{13}$ | 68 |
| RM-009 | 2 | Male | Intravenous | CAG-mNeonGreen, CAG-mRuby2, CAG-mTurquoise2 cocktail | 0.956 | $5 \times 10^{13}$ | 29 |
| RM-010 | 2 | Male | Intravenous | CAG-mNeonGreen, CAG-mRuby2, CAG-mTurquoise2 cocktail | 0.522 | $5 \times 10^{13}$ | 77 |

**All macaques in Supplementary tabe 2 were injected with CAP-Mac packaging the listed cargo.*

**Supplementary table 3: Green monkey information for individual characterization of CAP-Mac.**

| Green monkey ID | Age (days) | Sex | Route of administration | Capsid | Cargo | Weight at injection (kg) | Total dose (vg/kg) | Expression length (days) |
|---|---|---|---|---|---|---|---|---|
| C010 | 200 | Male | Intravenous | AAV.CAP-MAC | CAG-EGFP | 1.18 | 7.50E+13 | 36 |
| C017 | 172 | Male | Intravenous | AAV.CAP-MAC | CAG-EGFP | 1.32 | 7.60E+13 | 36 |
| C002 | 215 | Male | Intravenous | AAV9 | CAG-EGFP | 1.31 | 7.50E+13 | 36 |
| C016 | 178 | Male | Intravenous | AAV9 | CAG-EGFP | 1.02 | 7.60E+13 | 37 |

**Supplementary Fig. 1: Generating DNA library to produce round 1 and round 2 viral library.**

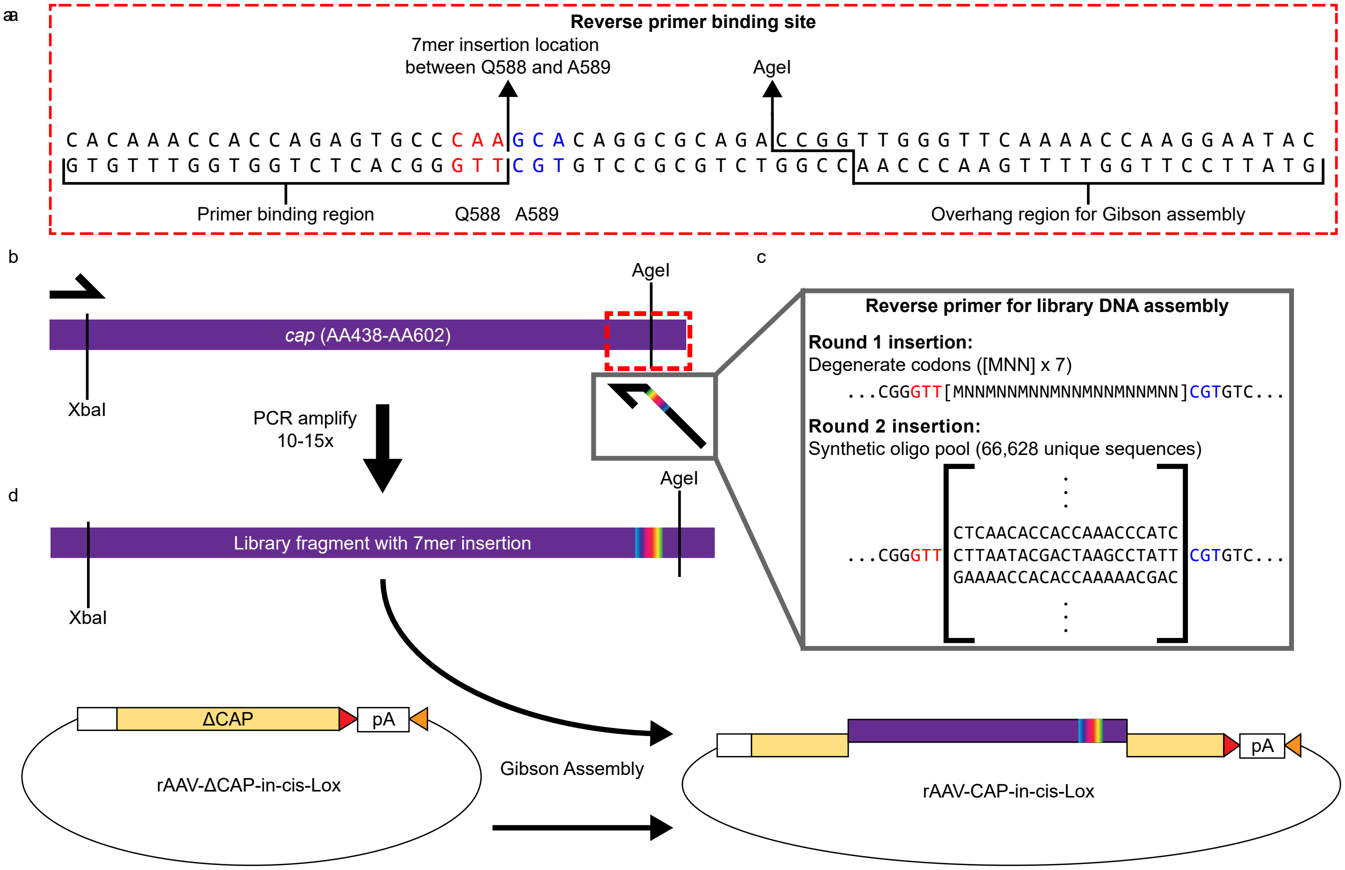


**Supplementary Fig. 1: Generating DNA library to produce round 1 and round 2 viral libraies. a,** Binding site on the AAV9 *cap* gene (proximal to the AA588 insertion location) for a reverse primer that is used to introduce diversity into the AAV9 genome. **b,** The reverse primer is used to generate a PCR fragment spanning the XbaI and AgeI section of *cap*, approximately AA438 to AA602. **c,** The reverse primer is designed such that it is homologous with the “Primer binding region” highlighted in (**a**), contains a 21 bp region that inserts a 7 amino acid sequence into the *cap* gene, and another homlogous sequence with *cap* that is necessary for subsequent Gibson assembly. For DNA assembly for round 1 selections, the reverse primer contains 21 degnerate codons ([MNN] x 7]). For round 2 selections, we used a synthetic oligo pool to specifically define each 21 bp sequence that we insert into the *cap* gene. **d,** After the PCR fragment containing our diverse region has been produced, we use Gibson assembly to create the final assembled library DNA.

**Supplementary Fig. 2: CAG-GCaMP7s expression after lumbar puncture administration of CAP-Mac.**

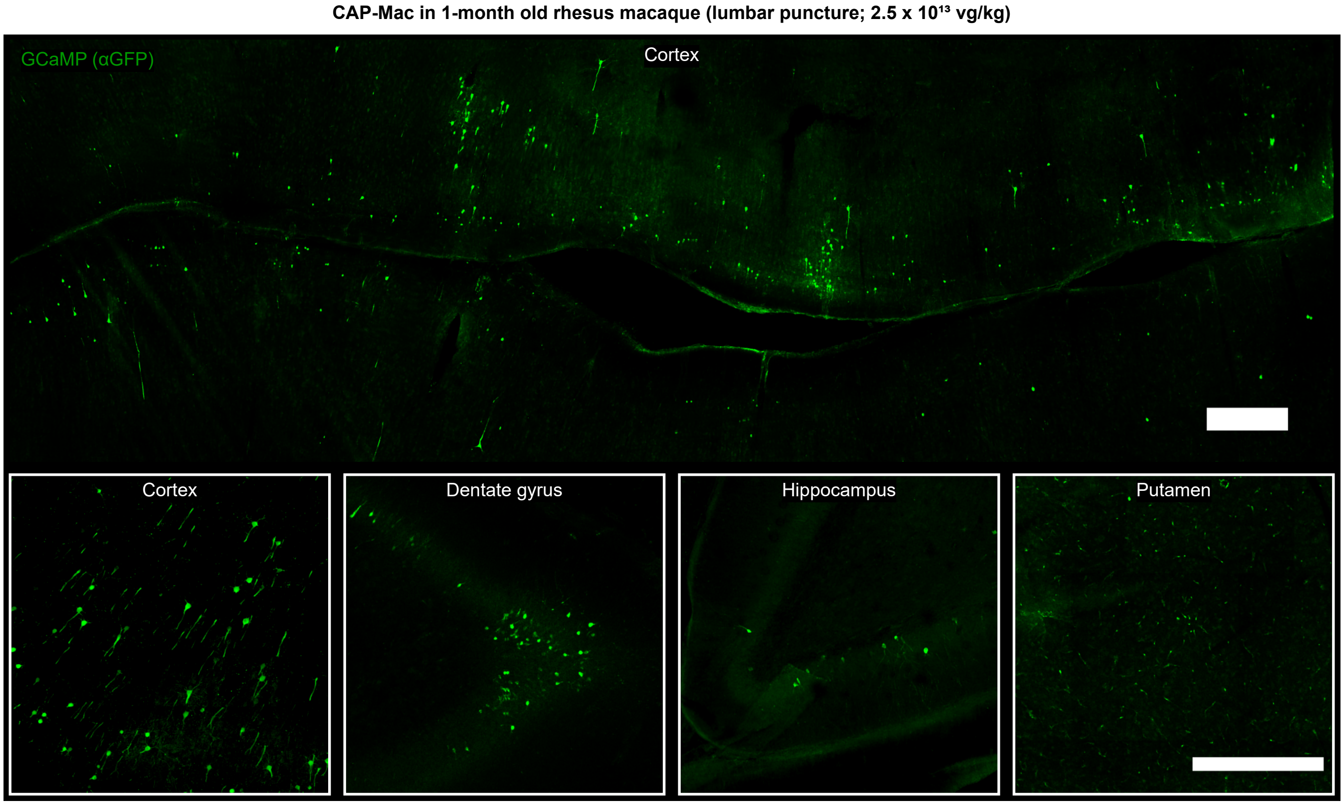


**Supplementary Fig. 2: CAG-GCaMP7s expression after lumbar puncture administration of CAP-Mac.** CAG-GCaMP7s expression detected in cortex, dentate gyrus, hippocampus, and putamen after staining with GFP antibody. All scalebars = 500 µm.

### Supplementary Fig. 3: CAG-XFP expression in non-brain tissue of Old World primates treated with AAV.

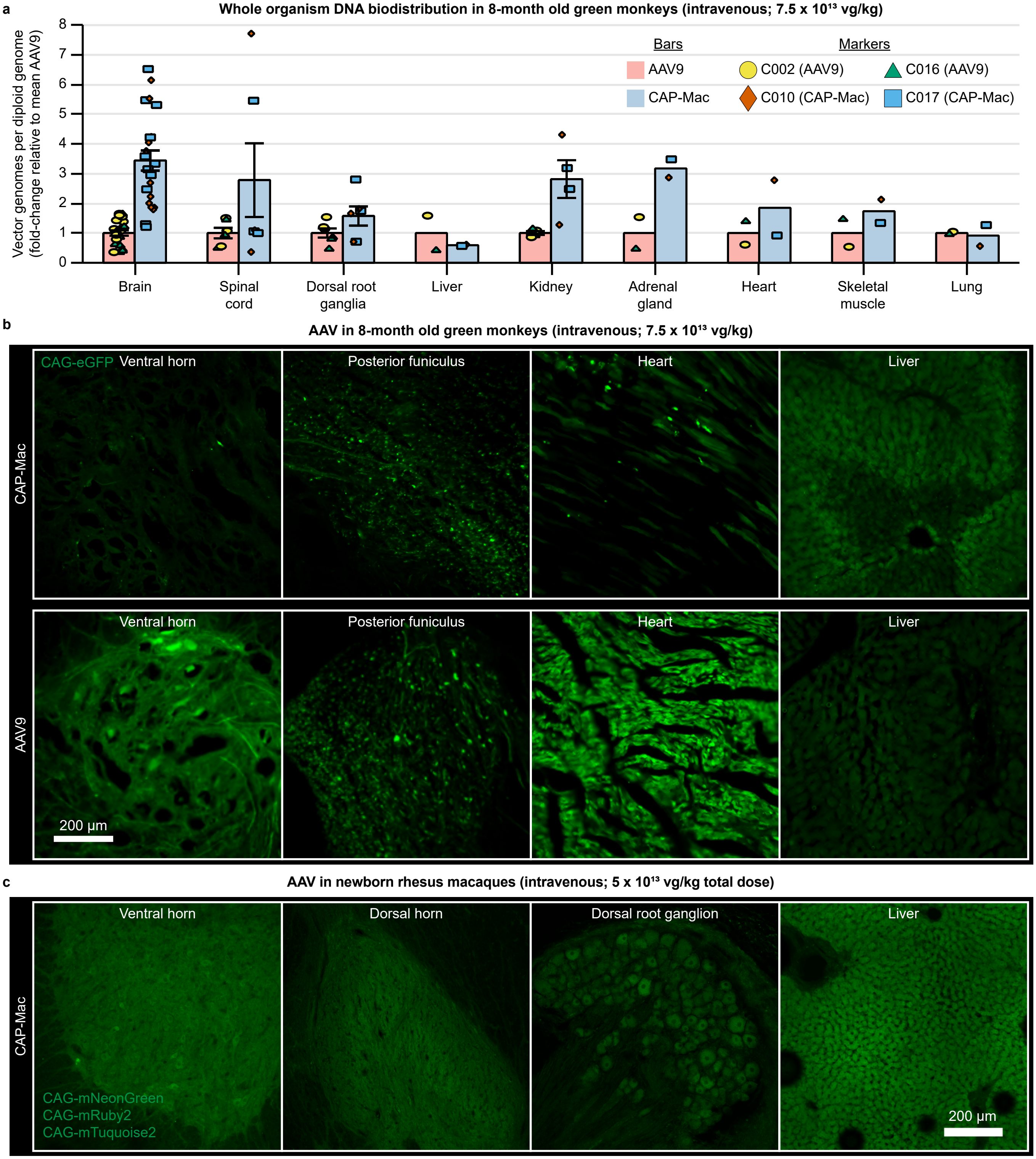


**Supplementary Fig. 3: CAG-XFP expression in non-brain tissue of Old World primates treated with AAV. a,** Vector genomes per diploid genome in 9 tissues from green monkeys treated with AAV9 and CAP-Mac, expressed as fold-change relative to mean AAV9. Red bars: mean AAV9 values. Blue bars: mean CAP-Mac values. Yellow circles: measurements from AAV9-treated monkey, C002. Green triangles: measurements from AAV9-treated monkey, C016. Orange diamonds: measurements from CAP-Mac-treated monkey, C010. Blue squares: measurements from CAP-Mac-treated monkey, C017. Each data point represents measured vector genomes per diploid genome in a piece of tissue from each condition. For the brain, spinal cord, dorsal root ganglia, and kidney, we sampled from 11, 3, 3, and 2 regions per animal, respectively. Mean ± s.e.m. shown (s.e.m. only calculated for samples with n > 2). **b**, CAG-eGFP expression in the spinal cord, heart, and liver of green monkeys after intravenous expression of either CAP-Mac (top) or AAV9 (bottom). **c**, CAG-XFP expression in the spinal cord, dorsal root ganglia, and liver of newborn rhesus macaque after intravenous administration of CAP-Mac packaging a cocktail of 3 CAG-XFPs. XFPs are pseudocolored identically.

**Supplementary Fig. 4: CAP-Mac in adult mice after intravenous and intracerebroventricular administration and in P0 mouse pups after intravenous administration.**

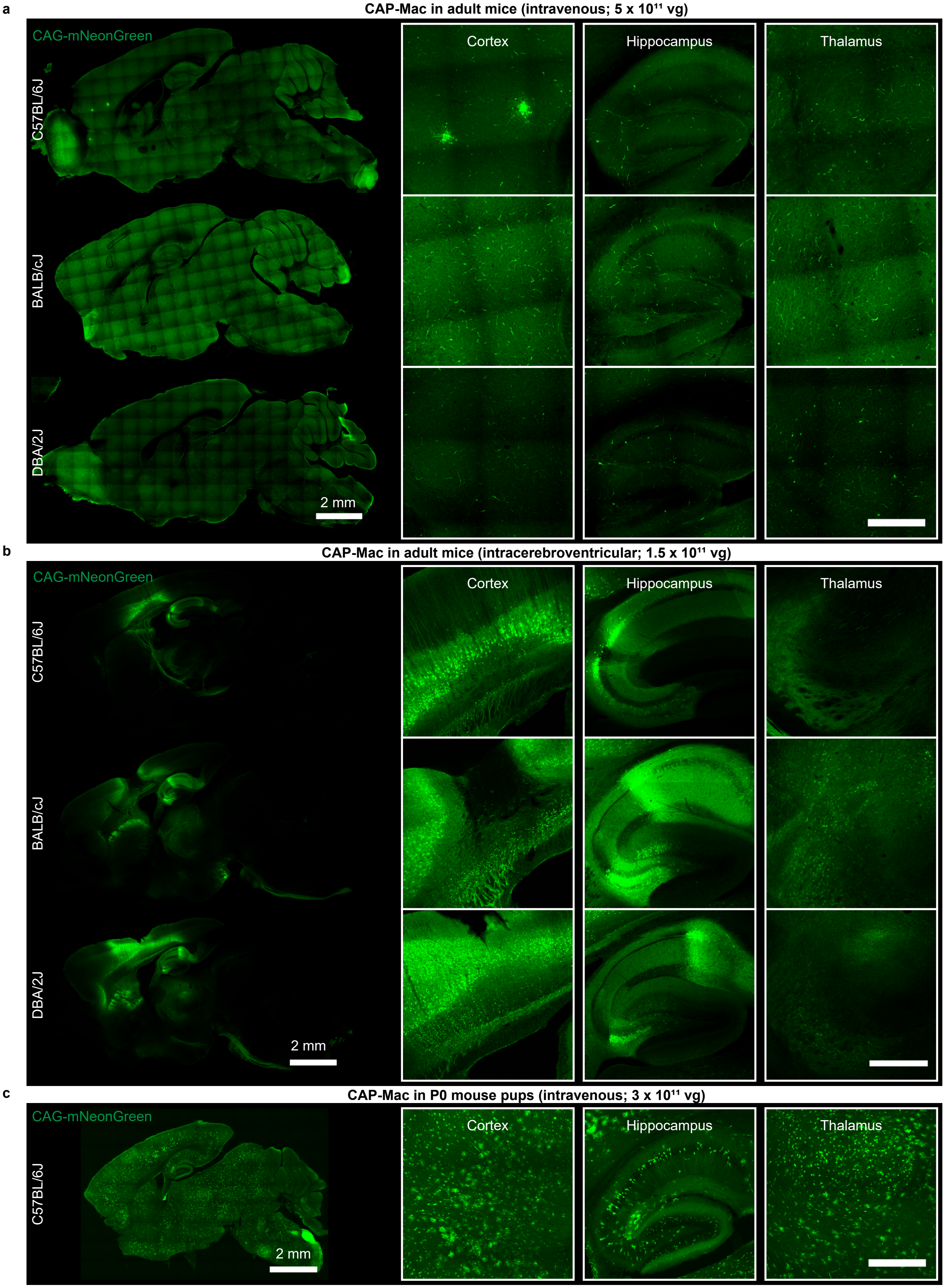


**Supplementary Fig. 4: CAP-Mac in adult mice after intravenous and intracerebroventricular administration and in P0 mouse pups after intravenous administration. a**, CAP-Mac after intravenous administration in C57BL/6J, BALB/cJ, and DBA/2J adult mice primarily transduces vasculature. **b**, CAP-Mac after intracerebroventricular administration in adult mice primarily transduces neurons. **c**, CAP-Mac in P0 C57BL/6J pups after intravenous administration transduces various cell-types, including neurons, astrocytes, and vasculature. All scalebars = 500 μm unless otherwise noted.
